# Natural Killer Cell Granule Protein (NKG7) promotes the development of vaccine-induced anti-fungal Th1 cells

**DOI:** 10.64898/2026.04.21.719793

**Authors:** Marcel Wüthrich, Cleison Ledesma Taira, Lucas dos Santos Dias, Xin He, Sarah Lichtenberger, Bruce S. Klein

## Abstract

CD4^+^ T cells that produce IFN-γ (T helper 1 [Th1] cells) chiefly mediate vaccine acquired resistance to fungal infections. However, the key regulators of the development of Th1 cells and acquired resistance to fungi are incompletely understood. Here, we report that Natural Killer Cell Granule Protein (NKG7) acts as an unappreciated regulator inducing anti-fungal Th1 cells to produce IFN-γ and converting plastic Th17 cells into polyfunctional memory Th1 cells.

**Author summary:** The mechanisms of vaccine resistance to fungi are incompletely understood. We identified a regulator that promotes the development of proinflammatory immune lymphocytes and fosters the conversion of one population of lymphocytes into multi-functional cells that produce several proinflammatory soluble factors that efficiently combat infectious diseases.

## INTRODUCTION

The role of IFNγ in antifungal resistance has been studied extensively. While IFNγ produced by CD4^+^ T cells promotes resistance to infection, too much IFNγ results in immunopathology and progressive infection [1]. Despite the accepted role of Th1 cells in antifungal resistance, protective strategies harnessing them is stunted by limited understanding of how best to promote their development.

The transcription factor T-bet, the primary transcription factor for development of Th1 cells, binds to the Natural Killer Cell Granule Protein (NKG7) promoter in Th1 cells, an understudied action that increases expression of both IFNγ and NKG7 [2]. NKG7-deficient CD4^+^ T cells respond poorly to IL-12 signaling, necessary for Th1 cell development [3]. We hypothesize that NKG7, first uncovered in natural killer cells and recently implicated in mediating Th1 cell differentiation, underpins the generation of antifungal immunity.

Despite an accepted role for Th1 cells in antifungal resistance [4-6], we and others have found a requisite role for Th17 cells in resistance to fungi [7]. In puzzling over how to reconcile seminal roles for both Th1 and Th17 subsets, we recently reported that antifungal Th17 cells convert into polyfunctional Th1 cells producing multiple cytokines including IFN-γ [8]. Therein, we used adjuvant formulations with glucopyranosyl lipid adjuvant (GLA) to enhance antifungal immunity. GLA induced “*plastic”* Th17 cells that convert into polyfunctional Th1 cells [8].

Here, we investigated the role of NKG7 in regulating development of antifungal Th1 cells and conversion of Th17 into Th1 cells during vaccine immunity. We report that: 1) NKG7 reporter mice vaccinated against fungi display increased IFN-γ expression: 2) Lack of NKG7 in global knockout mice or conditional mice lacking expression in CD4^+^ T cells result in blunted IFN-γ expression during Th1 cell differentiation and impaired antifungal resistance. 3) NKG7 regulates conversion of plastic Th17 cells into protective, multifunctional Th1 memory cells.

## RESULTS & DISCUSSION

### NKG7 as a correlate of antifungal Th1 cells

To identify unrecognized correlates of antifungal vaccine resistance, we compared the transcriptome of protective vs. non-protective CD4^+^ T cells [9]. In experimental blastomycosis, we have reported that C57BL6 mice vaccinated subcutaneously (SC) with *Blastomyces*-endoglucanase 2 (*Bl*-Eng2) acquire resistance [6], but those vaccinated intranasally (IN) do not. We extend those results here to include 129 and BALB/c mouse strains (**SFig. 1A+B**) to establish that the finding is not specific to a mouse genotype. We thus surmise that genes expressed by T cells from SC-but not IN-vaccinated mice are linked with vaccine resistance.

We compared *Bl*-Eng2-tetramer+ cells by scRNAseq in SC and IN vaccinated mice [9]. We observed that the percentage of tetramer+ T cells expressing IFNγ is higher in SC vaccinated mice than in IN vaccinated mice (**Fig. 1A**), underscoring IFNγ as a correlate of resistance [9]. Surprisingly, IL-17 expressing T cells were more abundant in IN vaccinated mice. We also found concordant expression between IFNγ and NKG7 in SC vaccinated mice but not IN vaccinated mice (**Fig. 1A**). Among16 distinct clusters of cells, NKG7 was expressed mainly in SC vaccinated mice by T cells that expressed IFN-γ, e.g. NK-like Th1 cells (nTh1 cells), Tis cells, and clusters 1, 3, 4 and 5 of the Th1/Th17 cell spectrum (**Fig. 1B**). NKG7 has been described as a regulator of IFNγ expression and resistance in experimental *Leishmania*s [3]. We hypothesized that NKG7 in CD4^+^ T cells regulates IFNγ expression and antifungal vaccine resistance. We tested this idea with different vaccine regimens in experimental blastomycosis, coccidioidomycosis and histoplasmosis.

### NKG7 expression in tetramer+ Th1 cells

To analyze NKG7 expression in IFN-γ producing T cells and vice versa, we created NKG7-reporter mice. We crossed NKG7-cre mice and mTmG reporter mice, vaccinated the mice with *Bl*-Eng2, and investigated co-expression of NKG7 and IFNγ. Tetramer+ T cells showed elevated NKG7, denoted by increased frequency of reporter+ cells vs. total CD4^+^ T cells (**Fig. 1C**). Likewise, the frequency of IFNγ+ T cells was increased among NKG7 reporter+ cells compared to reporter-T cells and total CD4^+^ T cells (**Fig. 1D**). Unvaccinated reporter mice did not generate tetramer+ T cells or cytokine+ T cells (**SFig. 1C**).

**Fig. 1:**
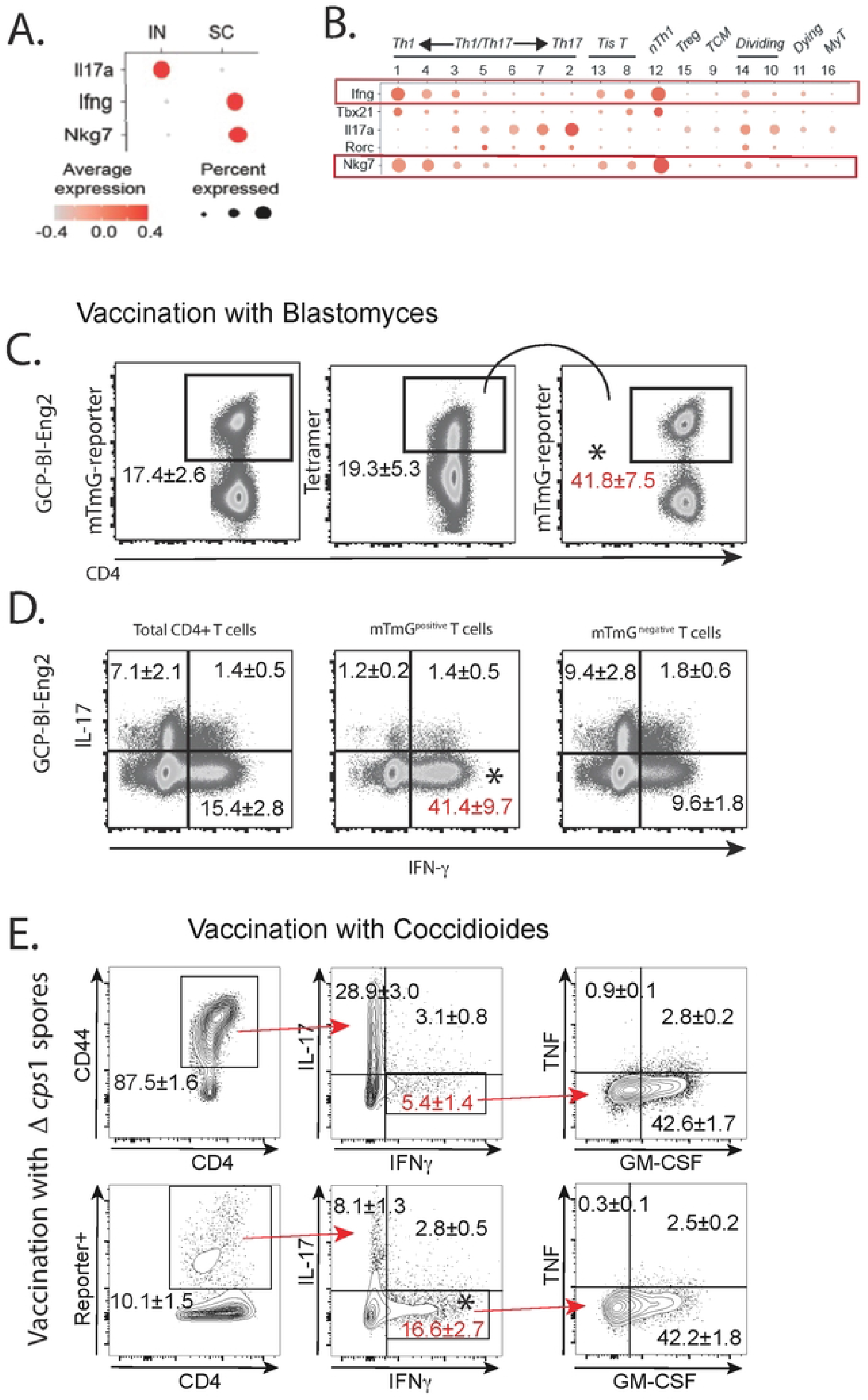
NKG7 expression is linked with vaccine resistance and IFNγ expression. (**A**) Transcripts associated with protection. (**B**) IFNγ and NKG7 expression mirror each other in T cell clusters. (**C+D**) NKG7 reporter mice were vaccinated with GCP-*Bl*-Eng2 twice two weeks apart. Two weeks after the boost, mice were infected with wild type *B. dermatitidis* yeast and lung T cells analyzed at day 4 post-infection. Lung T cells from GCP-*Bl*-Eng2 vaccinated mice were analyzed for the frequency of reporter, tetramer and cytokine positive cells. Dot plots are a concatenation of 5 mice/group, representing two independent experiments. The frequencies of cells are calculated as geometric means ± geometric SD. *p<0.05 vs. all other groups, Anova test. (**E**) NKG7 reporter mice were vaccinated with attenuated spores of *C. posadasii strain Δcps1* and rested for 6 months before challenge with wild-type arthroconidia. At day 6 post-infection lung T cells were stimulated *ex vivo* with *Δcps1* spores and analyzed for intracellular cytokine production. *p<0.05 vs. CD4^+^ CD44^+^ cells, Anova test.

We sought to exclude that our results were skewed by the vaccine antigen or fungal pathogen. To engage a larger, polyclonal T cell population and study the association between NKG7 and IFNγ in memory Th1 cells for other fungi, we used live vaccines against *Coccidioides posadasii* or *Histoplasma capsulatum*. We vaccinated NKG7-reporter mice with spores of attenuated *C. posadasii Δcps1* [10] or *H. capsulatum* yeast [11] since acquired resistance to both fungi requires Th1 immunity. Six months post-vaccination, we challenged reporter mice with corresponding wild type *Coccidioides* arthroconidia or live *Histoplasma* yeast to recall memory T cells to lung. In the *Coccidioides* model, NKG7+ reporter T cells expressed IFNγ at ≈3-fold higher frequency than total CD4^+^ T cells (**Fig. 1E + SFig. 1D**). Similarly, in the *Histoplasma* model, NKG7+ reporter T cells expressed IFNγ at ≈3-fold higher frequency than total CD4^+^ T cells (**SFig. 2**). Thus, IFNγ is linked with NKG7 expression in multiple models of fungal vaccination and among antigen specific T cells from narrow effector and large polyclonal memory populations.

### Requisite role of NKG7 in resistance and Th1 cell differentiation

To test whether NKG7 is required in acquired antifungal resistance, we vaccinated NKG7 global knockout mice and controls against blastomycosis with *Bl*-Eng2 and coccidiomycosis with *Δcps1* spores and measured lung CFU after infection. CFU values were several logs higher in vaccinated NKG7^-/-^ than controls (**Fig. 2A+B**).

We next investigated the role of NKG7 specifically in CD4^+^ T cells. We CRISPR-edited NKG7 in naïve, endogenous CD4^+^ T cells [12], which yielded >90% gene deletion (**Fig. 2C**). We adoptively transferred these NKG7-edited CD4^+^ T cells (NKG7^-negative^) or control edited (NKG7^-positive^) T cells into Rag1^-/-^ mice (lack endogenous T cells). We vaccinated recipients against blastomycosis with live attenuated yeast to engage polyclonal T cells [13, 14]. Among vaccine-induced CD4^+^ T cells recruited to lungs after challenge, editing of NKG7 (**Fig. 2D**) reduced IFNγ+ cells by 70%. Vaccinated mice that received NKG7^-negative^ T cells also died much faster after challenge than mice that received NKG7^-positive^ T cells (**Fig. 2E**). Remarkably, mice that received the NKG7^-negative^ T cells died as quickly as unvaccinated susceptible mice. NKG7^-negative^ recipients also showed significantly more lung CFU (**Fig. 2F**) than NKG7^-positive^ controls. These data support our premise that NGK7 regulates Th1 cell differentiation and is essential in antifungal vaccine resistance.

**Fig. 2:**
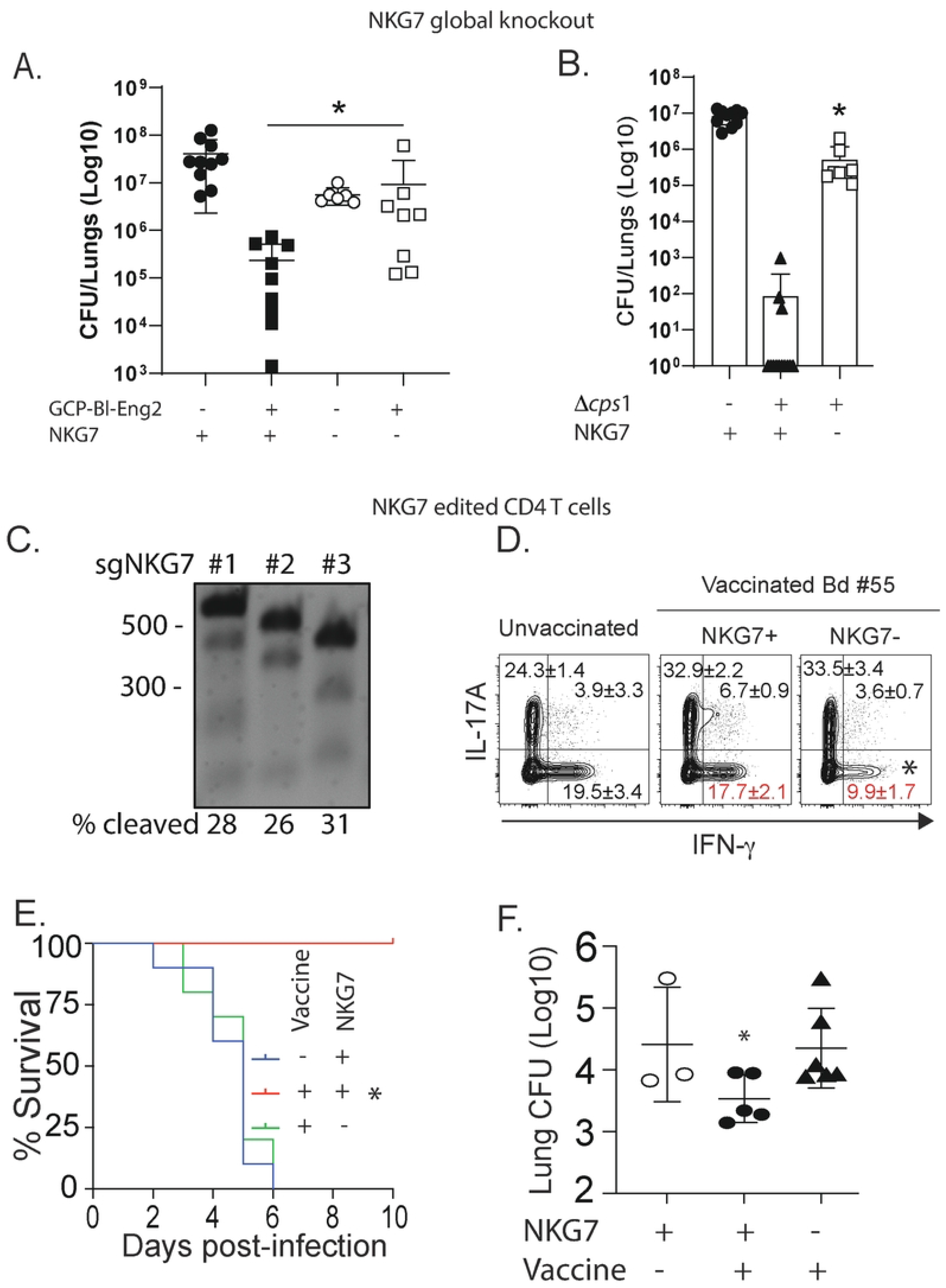
Role of NKG7 in vaccine resistance against blastomycosis and coccidiomycosis. Global NGG7^-/-^ mice were vaccinated with GCP-*Bl*-Eng2 or GCP-MSA as a control (**A**) or *Δcps1* spores (**B**) twice, two weeks apart. Two weeks after the boost, mice were infected with wild type *B. dermatitidis* yeast or *C. posadasii* arthroconidia and analyzed for lung CFU at the time controls appeared moribund *p<0.05, 2-tailed Mann-Whitney T test. (**C-F**) Impact of CRISPR editing NGK7 in T cells. (**C**) Naïve CD4+ T cells were enriched from naïve C57BL/6 mice and editing performed with three different sgRNAs combined targeting NKG7. The data show the genomic cleavage efficiency for each individual sgRNA: 28%, 26%, and 31%, respectively. The calculated combined cleavage efficiency is 63.3%, and the actual loss of the gene likely exceeds 80% of the transcript. (**D**) CRISPR-edited T cells were adoptively transferred into Rag1^- /-^ mice subsequently vaccinated with attenuated BAD1-null *B. dermatitidis* yeast. Three weeks later, mice were infected with wild type yeast and lung T cells analyzed for intracellular cytokine production at day 7 post-infection. (**E**) Survival after infection of vaccinated mice and controls and (**F**) Lung CFU. *p<0.05 vs. all other groups, Anova test.

In an orthogonal approach, we engineered conditional loss of NKG7 mice in endogenous T cells by crossing NKG7^flox/flox^ mice [3] with CD4-Cre mice. We vaccinated these mice (and NKG7^flox/flox^ controls) against *Coccidioides* infection with the protective antigen *Cp*-Eng2 as described [15] and gave tamoxifen before and throughout vaccination to ablate NKG7 expression in the endogenous T cells. To verify NKG7 ablation in conditional mice, we analyzed CD4^+^ T cells from the spleen of vaccinated and challenged mice. NKG7 protein expression was absent in conditional depleter mice compared to CD4-Cre-^negative^ controls (**Fig. 3A**). Moreover, the frequency of IFN_γ+_ T cells was sharply reduced in the mice that lacked NKG7 expression in CD4^+^ T cells compared to controls (**Fig. 3C+D**). Despite this defect in Th1 cell differentiation, the expansion of tetramer+ (**Fig. 3B**) and IL-17 producing T cells was similar in T cells lacking NKG7 compared to controls (**Fig. 3C+D**).

**Fig. 3:**
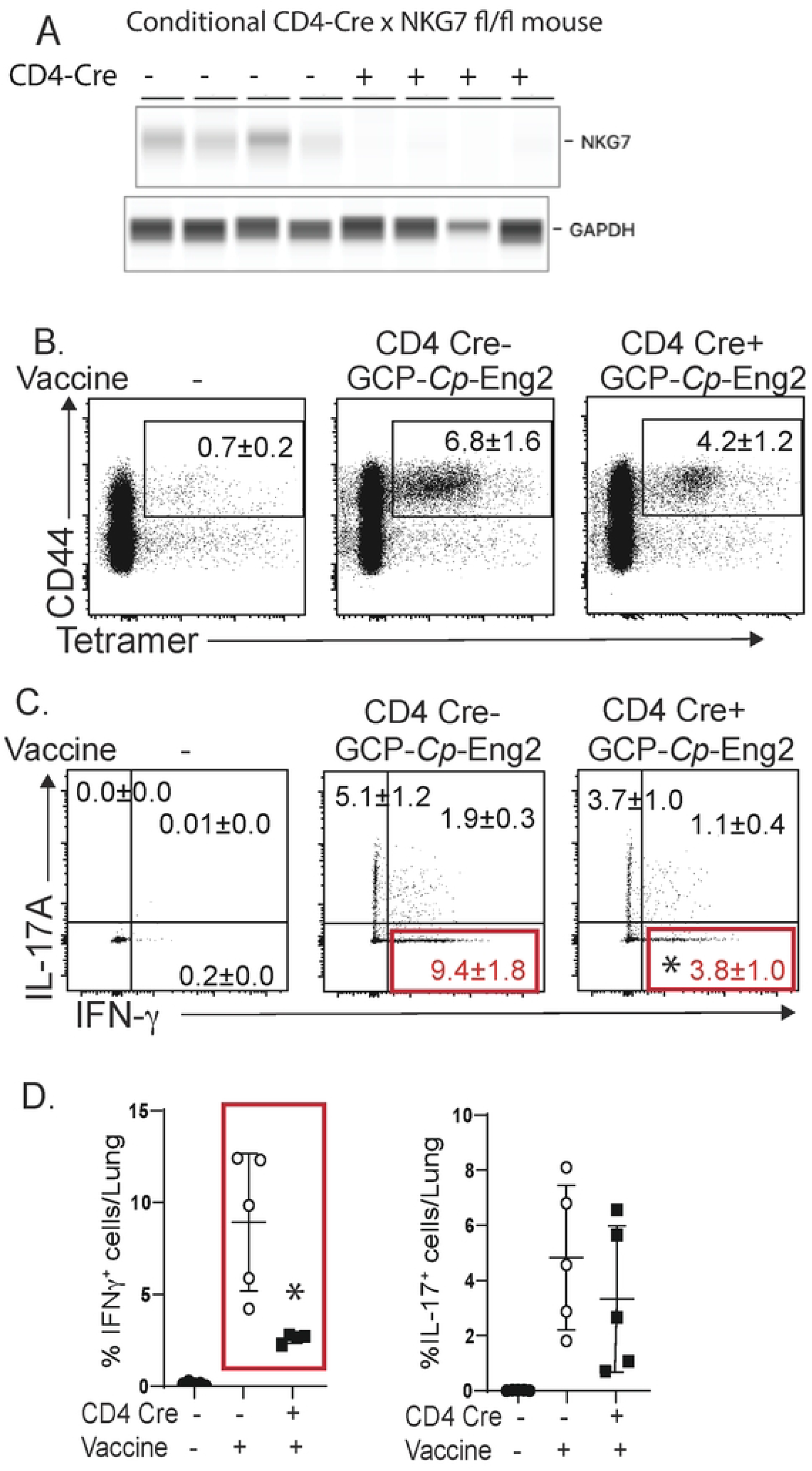
Role of NKG7 in CD4^+^ T cells for vaccine resistance against coccidioidomycosis. Mice were vaccinated against *C. posadasii* with GCP-*Cp*-Eng2. (**A**) Conditional CD4-Cre x NKG7^fl/fl^ and controls received tamoxifen prior to and during vaccination with GCP-*Cp*-Eng2 to ablate NKG7 in CD4^+^ T cells. At day 4 postinfection, NKG7 protein expression was determined by quantitative western blot in purified CD4^+^ T cells from the spleen. Lung T cells were analyzed for the presence of tetramer+ cells (**B**) and intracellular cytokine production (**C+D**).

These results support the idea that antigen-specific CD4^+^ T cells require NKG7 to differentiate into Th1 cells and mediate antifungal vaccine resistance. Our findings agree with the notion that NKG7 regulates lymphocyte function and downstream inflammation [3]. Depending on the context, NKG7 expression reportedly may enhance or blunt host response to infection. In experimental leishmaniasis, NKG7 is expressed by CD4^+^ T cells in infected and inflamed liver and spleen where it promotes expansion, recruitment and differentiation of protective Th1 cells [3]. Conversely, in experimental malaria, NKG7 mediates inflammation and recruitment of cytotoxic CD8^+^ T cells that damage brain vascular endothelium [3]. Since properly regulated inflammation enhances fungi killing, we conclude that NKG7 underpins acquired Th1 immunity needed to resist fungi.

### NKG7 promotes Th1 memory development with polyfunctional properties

We recently reported that adjuvant combinations including glucan chitin particles (GCP) with GLA drive the development of “plastic” Th17 cells that convert to polyfunctional Th1 memory T cells after vaccination and protect mice against experimental blastomycosis [8]. To investigate whether NKG7 regulates conversion of plastic Th17 cells into polyfunctional memory Th1 cells, we vaccinated conditional NKG7 mice with GCP-*Bl*-Eng2+GLA. We rested the mice for six months post-vaccination to allow memory cell formation and conversion of plastic Th17 into polyfunctional Th1 cells. We observed reduced frequencies of IFN_γ+_ T cells in mice with CD4^+^ T cells lacking NKG7 compared to controls (**Fig. 4A+B**), despite similar frequencies of tetramer+ cells in the groups. Additionally, 70% of the IFN-γ positive T cells from Cre^-negative^ control mice were polyfunctional and expressed GM-CSF and TNF, whereas only 56% of IFN_γ+_ T cells from Cre^-positive^ (NKG7-deficient) mice were polyfunctional (**Fig. 4B**). The frequency of IL-17 producing T cells conversely increased in NKG7-deficient mice compared to NKG7-sufficient controls (**Fig. 4B**). Thus, in the absence of NKG7, the ratio of IFNγ to IL-17 producing T cells became skewed toward IL-17 (**Fig. 4B**). Finally, lung CFU were significantly higher in conditional NKG7-deficient mice compared to NKG7-sufficient controls after infection (**Fig. 4C**), again indicating that NKG7 regulated expression of Th1 cell differentiation is required for antifungal vaccine resistance. Thus, NKG7 fosters both the differentiation of “straight” Th1 cells and also the conversion of plastic Th17 cells into polyfuntional memory Th1 cells.

**Fig. 4:**
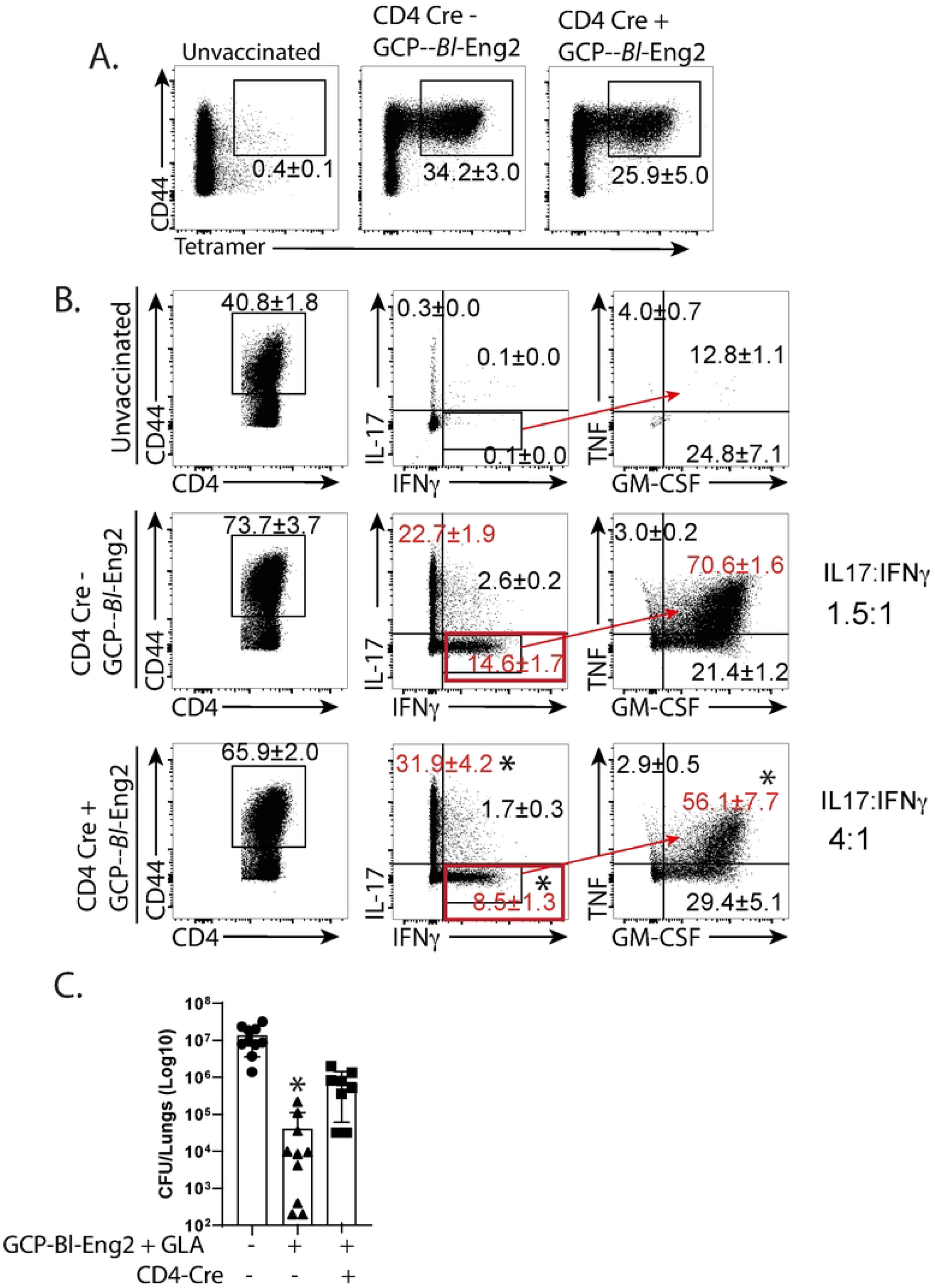
The role of NKG7 in CD4^+^ memory T cells in mice vaccinated against blastomycosis. Mice were vaccinated as in figure 1 with GCP-*Bl*-Eng2. Conditional CD4-Cre x NKG7 floxed mice received tamoxifen prior to and during vaccination to ablate NKG7 expression in CD4^+^ T cells. At day 4 postinfection lung T cells were analyzed for the presence of tetramer+ cells (**A**) and intracellular cytokine production (**B**). *p<0.05 vs. Cre-mice, Anova test. (**C**) Lung CFU 14 days after infection. *p<0.05 vs. all other groups, 2-tailed Mann-Whitney T test.

In summary, the role of NKG7 in infectious disease is understudied and the regulation of NKG7 in antifungal Th1 immunity remains important given the centrality of IFNγ and Th1 cells in orchestrating antifungal defense. NKG7, first identified in natural killer cells and cytotoxic T cells [16], has been studied chiefly in these cells where it mediates cytotoxicity and degranulation by association with perforin and granzyme release [3]. NKG7 also restrains mTORC1 activity in CD8^+^ T cells, supporting T cell durability and memory formation - keys to long term immunity [17].

The pivotal role of IFNγ in shaping immunity to fungi and other microbes make further work on NKG7 germane for harnessing the immune system for therapeutic benefit. Important questions remain about whether Th17 lineage memory cells are superior to Th1 lineage cells in protection against fungi and about the transcriptional profiles of these two Th1 cell lineages. By elucidating a key regulator of acquired antifungal resistance, such work will advance design of immune-based, host-directed therapy.

## METHODS

Animal studies adhered to protocol M005891 approved by UW-Madison’s IACUC. For additional methods, see <u>S1 Text</u>

## Acknowledgements

We thank Dr. Chris Engwerda from QIMR, Berghofer Medical Research Institute in Brisbane, Australia for generously providing NGK7-floxed, NKG7 knockout and NKG7-cre mice and his advice. The work was supported by NIH grants R01AI93553 (MW), R01AI040996 (BK/MW), R01 AI168370 (BK), U01 AI124299 (BK), R37 AI035681 (BK) and R24AI192252. XH is a Cancer Research Institute Irvington Fellow supported by the Cancer Research Institute (CRI4476). Flow samples were processed at the University of Wisconsin Carbone Cancer Center (UWCCC) Flow Core Facility on a BD LSR Fortessa that was purchased with the NIH shared instrumentation grant 1S100OD018202-01 and University of Wisconsin Carbone Cancer Center Support grant P30 CA014520.

## FIGURE LEGENDS

**SFig. 1: Resistance against blastomycosis in inbred strains of mice**. Balb/C and 129 mice received vaccine subcutaneously (SC) or intranasally (IN) with GCP-*Bl*-Eng2 twice, two weeks apart and were rested for two weeks before challenge with wild type strain 26199 yeast of *B. dermatitidis*. (**A**) Lung CFU were measured when controls were moribund. *p<0.05, 2-tailed Mann-Whitney T test. (**B**) Number of IFN_γ+_ CD4^+^ T cells at day 4 post-infection. (**C**) Vaccination of NKG7 reporter mice with GCP-MSA as a control and *ex vivo* recall with *Bl*-Eng2 peptide. (**D**) NKG7 reporter mice were vaccinated SC with *Δcps1* spores and challenged six months later with wild type *C. posadasii* arthroconidia; 6 days later, lung T cells were recalled with Pra14 peptide in the presence of anti-CD28 mAb and analyzed for intracellular cytokines. *p<0.05 vs. CD4^+^ CD44^+^ cells, Anova test.

**SFig. 2: Response of NKG7 reporter mice to vaccination against histoplasmosis**. Mice were vaccinated subcutaneously with live *Histoplasma* yeast and 6 months later challenged with a sublethal dose of 10^6^ wild type *Histoplasma* yeast. At day 6 post-infection, lung T cells were recalled *ex vivo* with cell wall/membrane (CW/M) extract or heat killed *Histoplasma* yeast and analyzed for intracellular cytokines. *p<0.05 vs. CD4^+^ CD44^+^ cells, Anova test.

## List of supplementary materials

S1 text file: Materials and Methods

S2 excel file: Raw data used to generate graphs will be submitted with the revision Supplementary Figure 1 to main Figure 1

Supplementary Figure 2 to main Figure 1

